# Memory-like B cells emerging from germinal centres recycle through the subcapsular sinus

**DOI:** 10.1101/2020.12.08.415828

**Authors:** Yang Zhang, Laura Garcia-Ibanez, Carolin Ulbricht, Laurence S C Lok, Thomas W Dennison, John R Ferdinand, Jennifer Mueller-Winkler, Cameron J M Burnett, Juan C Yam-Puc, Lingling Zhang, Geoffrey Brown, Victor L J Tybulewicz, Antal Rot, Anja E Hauser, Menna R Clatworthy, Kai-Michael Toellner

## Abstract

Infection or vaccination leads to the development of germinal centers (GCs) where B cells evolve high affinity antigen receptors, eventually producing antibody-forming plasma cells or memory B cells. We followed the migratory pathways of B cells emerging from germinal centers (B_EM_) and found that many migrated into the lymph node subcapsular sinus (SCS) guided by sphingosine-1-phosphate (S1P). From there, B cells may exit the lymph node to enter distant tissues. Some B_EM_ cells interacted with and took up antigen from SCS macrophages, followed by CCL21-guided return towards the GC. Disruption of local CCL21 gradients inhibited the recycling of B_EM_ cells and resulted in less efficient adaption to antigenic variation. Our findings suggest that the recycling of B_EM_ cells, that transport antigen and that contain the genetic code for B cell receptor variants, may support affinity maturation to antigenic drift.

---

The hallmark of adaptive immunity is memory, which is mediated by the expansion and long-term survival of antigen-specific lymphocytes, affinity maturation of B lymphocytes, and the long-term production of neutralizing antibody. Affinity maturation of B cells occurs via molecular evolution in germinal centers (GCs) (Zhang et al., 2016). This involves cycles of B cell proliferation and the mutation of B cell receptor genes, followed by natural selection of B cells expressing the highest affinity B cell receptors. The output of the GC reaction is high affinity antibody-producing plasma cells and memory B cells, both providing long-term immunity (Weisel et al., 2016; Yoshida et al., 2010; Zhang et al., 2018).

Plasma cells can be very long-lived (Landsverk et al., 2017), as are memory B cells (Gitlin et al., 2016; Pape et al., 2018). Interestingly, the affinity-dependent selection of memory B cells in the GC is less stringent than that seen for plasma cells, resulting in a highly variable pool of antigen-specific cells (Suan et al., 2017). As long-term immunity can be provided by long-lived plasma cells, the advantage of a low quality B cell output from the GC is not immediately obvious (Zinkernagel, 2018). However, their high variability may provide a pool of cells with the potential to protect against pathogen variants. Memory B cells can sense specific antigen, rapidly enter secondary responses, immediately present antigen to memory T cells (Ise et al., 2014; Jelcic et al., 2018), and generate new plasma cells within days (Mesin et al., 2020; Moran et al., 2018; Toellner et al., 1996).

Lymph nodes are important sites for the initiation of the adaptive immune response. They represent a platform where immunological information is sequestered and exchanged. Resident cells, including B cells, occupy distinct anatomical niches, and their movement between different areas of the lymph node is required for the progression of a GC reaction (Yi et al., 2012). One important structure in this regard is the subcapsular sinus (SCS), the primary area into which tissue derived lymph fluid drains, bringing antigens and pathogens. The SCS houses a subset of CD169^+^ macrophages that are specialized for antigen acquisition and pathogen defense (Moseman et al., 2012) and shuttle antigen to naïve and memory B cells (Arnon et al., 2011; Carrasco and Batista, 2007; Junt et al., 2007; Moran et al., 2018).

## Results

### Appearance of memory like B cells entering the subcapsular sinus guided by S1PR

In order to track the migration of antigen-specific B cells and plasma cells as they emerged from primary GCs in draining lymph nodes (drLN) following immunization, we adoptively transferred 4-hydroxynitrohpenyl (NP)-specific B cells from B1-8i mice (Sonoda et al., 1997), which express eGFP under the control of the Prdm1 promoter (Kallies et al., 2004) (labelling plasmablasts and plasma cells with eGFP) and Cdt1-mKO2 hybrid protein (Sakaue-Sawano et al., 2008) (labelling cells in G0/G1 phase of cell cycle with mKO2), and immunized with NP coupled to the carrier protein chicken gamma globulin (CGG). As we previously described (Zhang et al., 2018), plasmablasts emerged from the interface between the GC dark zone and T cell areas (Fig. 1A). Large numbers of antigen-specific B cells were located in the outer follicle surrounding GCs, typically close to the LN SCS (Fig. 1A). Cdt1-mKO2 labelling of these B cells suggests that they were recently activated B cells that emerged from adjacent GCs (B_EM_). This is reminiscent of historical observations describing the accumulation of marginal zone-like memory B cells under the SCS (Liu et al., 1988; Stein et al., 1980), and recent descriptions of switched memory B cells in follicles around GCs and under the SCS (Aiba et al., 2010; Moran et al., 2018).

**Fig 1:**
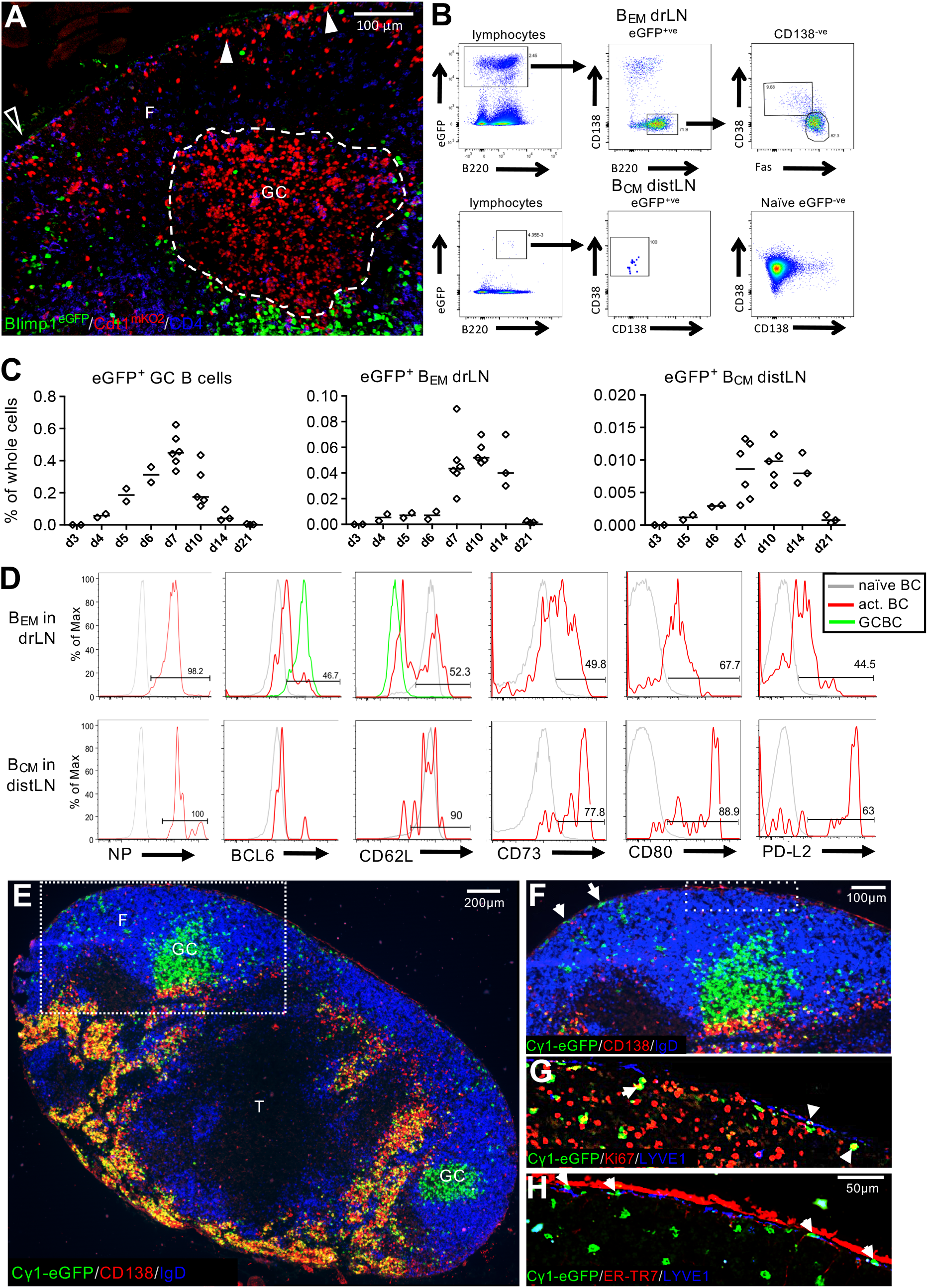
Appearance of antigen-activated memory-like B cells in drLN and distLN. **A**) Location of B1-8i/k^-/-^/Blimp1^GFP^/Cdt1m^KO2^ B cells in drLN 6 d after immunization. Antigen-specific B cells in the G0/G1 phase of cell cycle (red) inside the GC (dashed line) and in the follicle (F) close to the SCS (arrow heads). Interfollicular region (open triangle). Blimp-1^+^ PCs are eGFP (green). Hoechst33342-labelled naïve T cells (blue). Scale bar: 100 μm. **B**) Gating of eGFP^+^ B_EM_ and B_CM_ in drLN and distLN 8 d after immunization of recipients of NP-specific Cγ1Cre QM mTmG B cells. **C)** Kinetic of eGFP^+^ B cell appearance in drLN and distLN. Data merged from two independent experiments. **D)** Memory B cell typical markers on B_EM_ and B_CM_ in drLN and distLN. **E)** drLN of recipient of QM Cγ1Cre mTmG cells 8 d after immunization. T zone (T) **F)** Enlargement of box in E) showing the eGFP^+^ B_EM_ close to the subcapsular sinus, **G)** Ki-67 expression in B_EM_, **H)** same area showing B_EM_ location below the LYVE1^+^ ER-TR7^-ve^ SCS floor endothelium and inside the SCS (arrowheads).

In order to identify GC-derived B_EM_ in drLNs and their migration to non-reactive distant lymphoid tissues (distLNs), we used the well-established Cγ1-Cre reporter strain, which induces constitutive expression of GFP in B cells after T cell-dependent activation, which includes GC-lineage B cells (Casola et al., 2006). We crossed these with mice expressing B cell receptors specific for the hapten 4-hydroxynitrophenyl (NP) (Cascalho et al., 1996; Marshall et al., 2011) and a Cre-inducible eGFP reporter (QM Cγ1Cre mTmG mice) (Casola et al., 2006; Muzumdar et al., 2007). We immunized wild type (WT) host mice that had received a small number of antigen-specific B cells from QM Cγ1Cre mTmG. We observed that GC B cells (eGFP^+^NP^+^CD38^low^Fas^+^) were detectable from 4 d after immunization with maximum numbers seen at 6 - 10 d (Fig. 1 B-D). Within a day of GCs reaching maximum size, there was the emergence of a population of B cells that were eGFP^+^, NP-binding, CD38^high^, Fas^int^, CD138^-^, Bcl6^low^ (Fig. 1B) in the drLN (Fig. 1C, D). These cells also started to express markers associated with memory B cells such as CD73, CD80 and PD-L2 (Weisel et al., 2016). At the same time, antigen-activated B cells were observed in distant lymphoid tissues (Fig1 B – D). These eGFP^+^ circulating memory B cells (B_CM_) were confirmed to be antigen-specific, expressed CD62L at similar levels to naïve B cells, and high levels of CD73, CD80, and PDL2 (Fig. 1D). This suggests that the presence of B_EM_ close to the SCS at the peak of the GC response is related to emigration of antigen-activated B cells from the drLN through the SCS, generating systemic cellular B cell immunity.

Further immunohistological examination of drLNs around the peak of B_EM_ migration (Fig. 1 E-H) showed that the eGFP^+^ B_EM_ in B cell follicles surrounding the GC were still in cell cycle (Fig. 1G). Staining with Lyve-1 and ER-TR7, to identify the SCS floor and ceiling respectively, showed that indeed some B_EM_ had moved into the SCS (Fig. 1H). These data suggest that B_EM_ move from the GC into the SCS, from where they may join the efferent lymph flow (Girard et al., 2012), leaving the drLN to disseminate via blood into distant lymphoid tissues.

Intravital imaging of drLN of Cγ1Cre mTmG mice confirmed that a large number of eGFP^+^ B_EM_ had actively migrated between the GC and the SCS (Fig. 2A, suppl. movie 1,2). B_EM_ entered the SCS lumen (Fig. 2B), where some moved along the SCS (Fig. 2C) presumably migrating towards efferent lymphatics. Surprisingly, some B_EM_, after a short pause around macrophages in the SCS, re-entered the LN follicles through the SCS floor and migrated back towards the GC (Fig. 2B, D).

**Fig 2:**
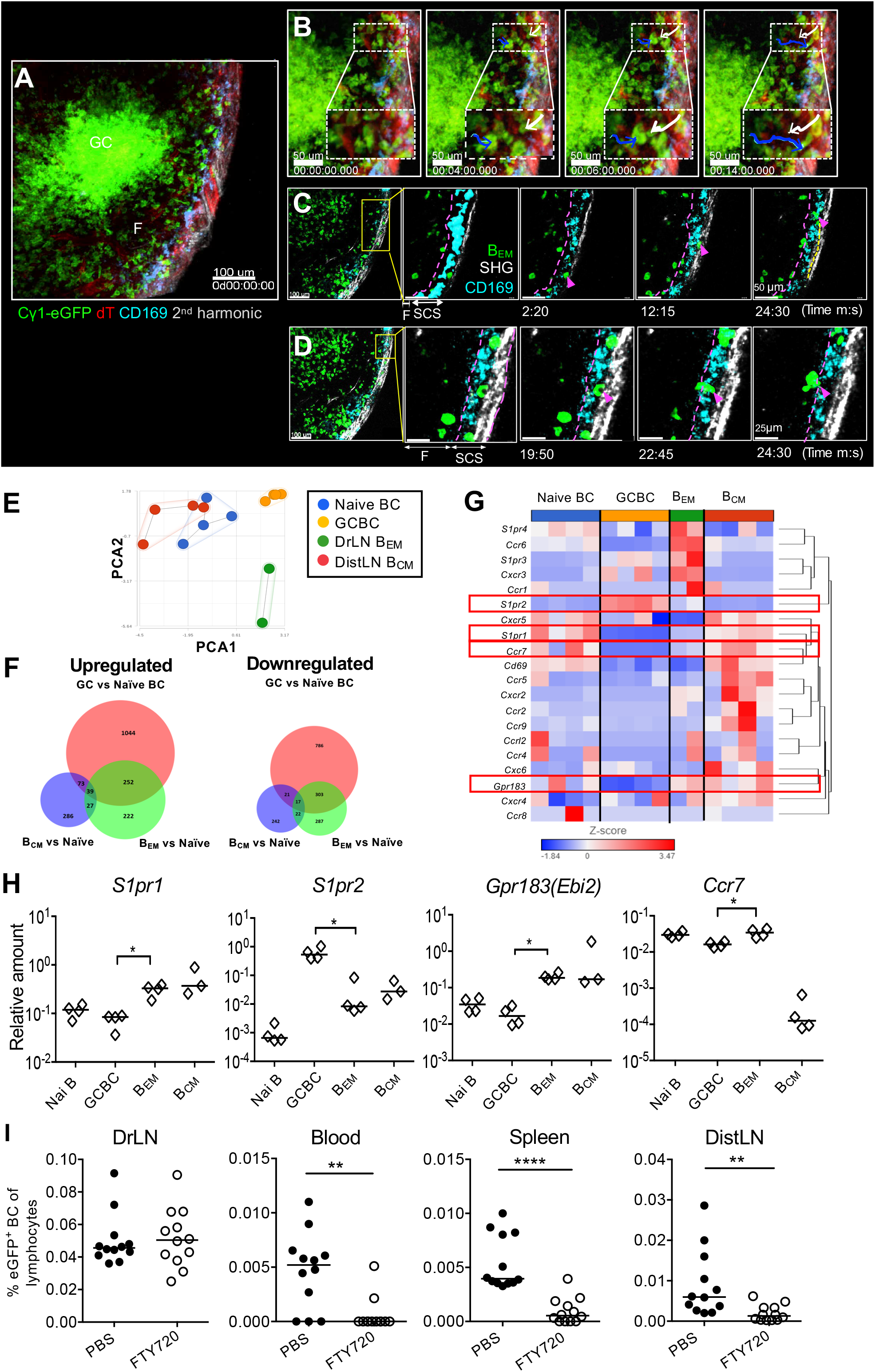
B_EM_ movement in the drLN. **A)** Intravital observation of popliteal lymph node from Cγ1Cre mTmG mice 8 d after NP-CGG foot immunization (see suppl. movie 1). **B)** Still images showing eGFP^+^ B_EM_ entering the SCS (blue arrow) and reentering the lymph node follicle from the SCS (white arrow). **C)** Images showing a eGFP^+ve^ B cells migrating along the SCS. **D**) Images showing a B_EM_ reentering the lymph node follicle. **E-G)** RNAseq data from B cell populations sorted from drLN 8 d after immunization of receipients of Cγ1-Cre QM mTmG B cells. **E)** Principle component analysis of global gene expression in sorted populations, **F)** Numbers of differentially expressed genes. **G)** Heat map of selected chemotactic receptors among mRNA. Genes were hierarchically clustered by Euclidean distance measure. Data are from two independent experiments with 4 samples (each sample from pooled popliteal lymph nodes from 3 mice). **H)** *S1pr1, S1pr2, Ebi2*, and *Ccr7* mRNA expression analyzed by qRT-PCR. Each diamond represents pooled lymph nodes from 4 mice. All values are relative to *B2m* mRNA. Two-tailed Mann-Whitney test, *: p<0.05. Data representative of four independent experiments. **I**): B_EM_ in drLN and B_CM_ in blood, spleen, and distLN 8 d after immunization. Mice received FTY720 over 2 d before sacrifice. Each dot represents one mouse, data merged from three independent experiments. Two-tailed Mann-Whitney test, **: p<0.01, ****: p<0.0001.

To examine the factors that regulate the migration of B_EM_ from GCs, we performed RNASeq analysis of FACS sorted eGFP^+^ B_EM_ in LNs comparing them to naïve B cells, GC B cells, and B_CM_ from distLNs. Principle component analysis of all genes expressed by these four subsets of cells confirmed a close relationship of eGFP^+^ B_EM_ with GC B cells, whereas eGFP^+^ B_CM_ in distLN are much closer to naïve B cells (Fig. 2E). This was also evident in the number of individual genes differentially expressed, with a greater number of genes differentially expressed regarding the transition from naïve to GC B cells, and a larger overlap in genes coexpressed in GC B cells and B_EM_ (Fig. 2F). Analysis of migratory receptors during the transition from GC B cells into B_EM_ by qRT-PCR revealed a loss of expression chemokine receptors known to be associated with B cells location in the GC (*Cxcr5, Cxcr4, S1pr2*) (Green and Cyster, 2012), and increased expression of the receptors *S1pr1, S1pr3, S1pr4, Ebi2, Cxcr3, Ccr6*, and *Ccr7* (Fig. 3G, H). CCR6, EBI2, and CXCR3 are known to be expressed on memory B cells (Stoler-Barak et al., 2019; Suan et al., 2017). Blockade or deletion of these receptors, however, did not lead to a noticeable change in the appearance of B_CM_ in distLNs. S1P receptors, particularly S1PR1 and S1PR2, are known to direct the location of B cells in the follicle center and their emigration into lymph vessels (Green and Cyster, 2012). *In vivo* S1PR blockade using FTY720 led to a dramatic reduction of B_CM_ in blood and distLNs (Fig 3I), while there was no noticeable effect on the numbers of other lymphocytes in distant lymphoid tissues. This suggests that S1PR guides memory B cell migration into the SCS and to lymphatic vessels.

**Fig 3:**
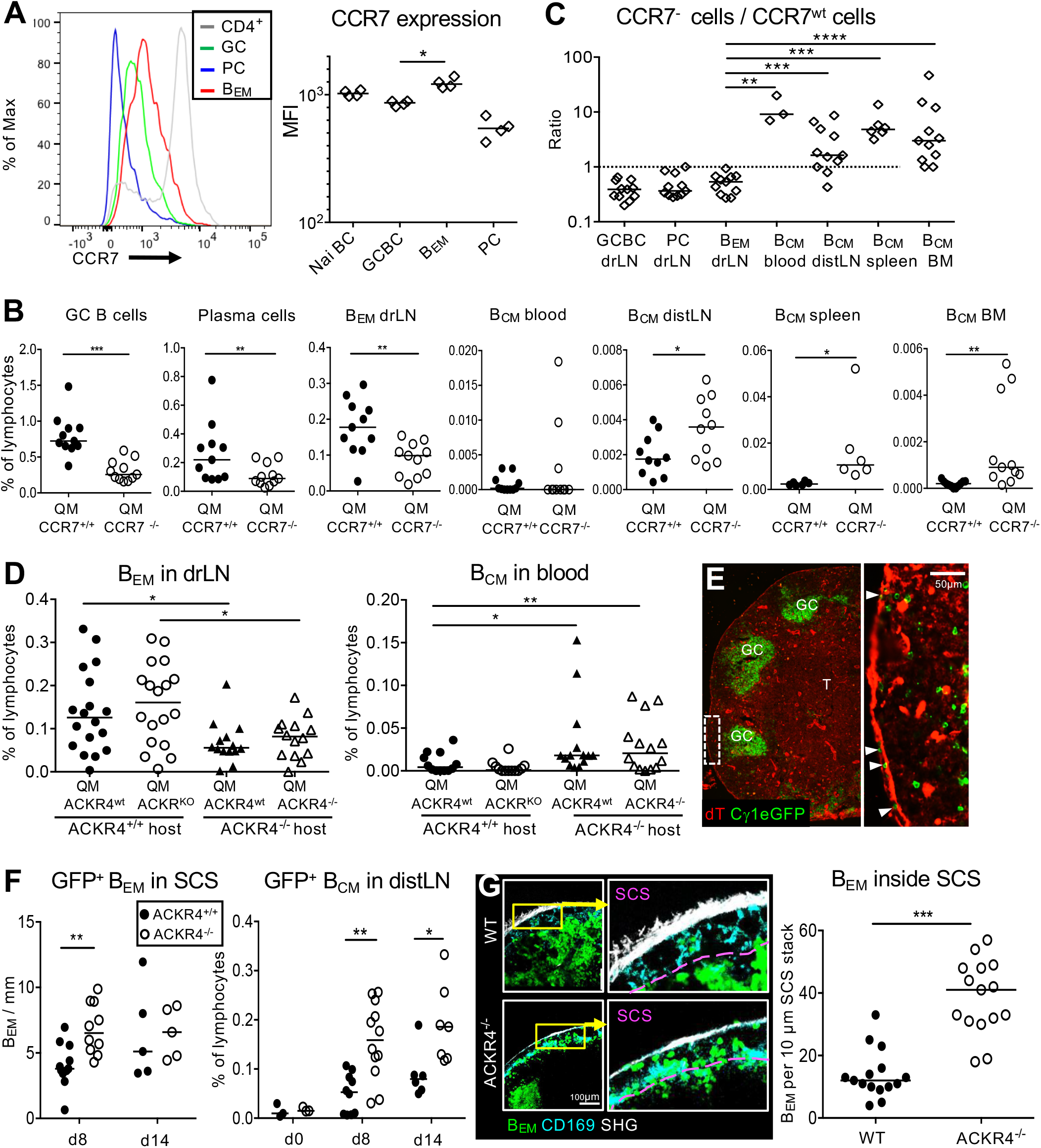
CCL19/21-dependent migration of memory B cells. **A)** CCR7 expression on different lymphocyte subsets measured by flow cytometry. **B)** GC B cells, plasma cells (PC), B_EM_ from drLN 8 d after foot immunization of recipients of a mix of QM CCR7^WT^ and QM CCR7^-/-^ B cells. B_CM_ in blood, distLN, spleen, and bone marrow (BM). **C)** Ratio of CCR7^+/+^ to CCR7^-/-^ eGFP^+^ QM B cells in different tissues. Each symbol represents one animal. Data are from two independent experiments with 5-6 mice. Different hosts compared using Wilcoxon test *p<0.05; **p<0.01; ***p<0.001. **D)** Frequency of ACKR4^+/+^ and ACKR4^-/-^ B_EM_ or B_CM_ in drLN or blood of ACKR4^WT^ and ACKR4^-/-^ hosts. Data from 3 independent experiments with 4-5 mice each. Mann-Whitney test, *p<0.05; **p<0.01. **E)** drLN from an Cγ1-Cre mTmG mouse 8 d after immunization. Arrows: eGFP^+^ B_EM_ inside the SCS. **F)** eGFP^+^ B_EM_ in the SCS of Cγ1Cre mTmG in drLN and B_CM_ in distLN in ACKR4^+/+^ or ACKR4^-/-^ mice. **G)** Intravital observation of B_EM_ entry into SCS in C*γ*1Cre mTmG ACKR4^+/+^ or ACRK4^-/-^ drLN. Quantitation of intravital eGFP^+^ B_EM_ in SCS of drLN. Each dot represents cell count per field of view of a 10 μm Z stack. Pooled data from imaging of 3 ACKR4^+/+^ and 2 ACKR4^-/-^ popliteal LNs. Unpaired t-test, \*\*\**p* < 0.001.

### CCR7 dependent recycling of B_EM_

The intravital imaging we performed (Fig. 2A-D) showed that many B_EM_ after entering the SCS returned to the follicles. Dendritic cells (DC), arriving in the SCS from afferent lymph migrate into the lymph node guided by local CCL21 gradients that are sensed by CCR7 on DC (Ulvmar et al., 2014). As B cells upregulate CCR7 during the transition from GC B cell to B_EM_ (Fig. 2H, 3A), we hypothesized that a local CCL21 gradient might have a similar role for B_EM_ return into the drLN, as it has for DCs. In order to test this, QM CCR7^+^ mT^+^ and QM CCR7^ko^ eYFP^+^ B cells were co-transferred into WT mice and their migration assessed after immunization. CCR7 is required for the initial activation of naïve B cells, enabling B cell migration into T cell zones (Okada et al., 2005; Reif et al., 2002). Therefore, CCR7-deficient B cells were underrepresented in activated B cell populations (antigen-specific GC B cells, plasma cells, and B_EM_) in the drLN (Fig. 3B). Despite this, there was an increase in *Ccr7*^-/-^ B_CM_ in blood, distLNs, spleen, and bone marrow (Fig. 4B, C). This is compatible with a role for CCL21 in orchestrating B_EM_ re-entry into the follicle from the SCS. Without cues sensed by CCR7, B_EM_ are unable to move back from the SCS into the LN parenchyma and therefore appeared in larger numbers in blood and distant lymphoid tissues.

**Fig 4:**
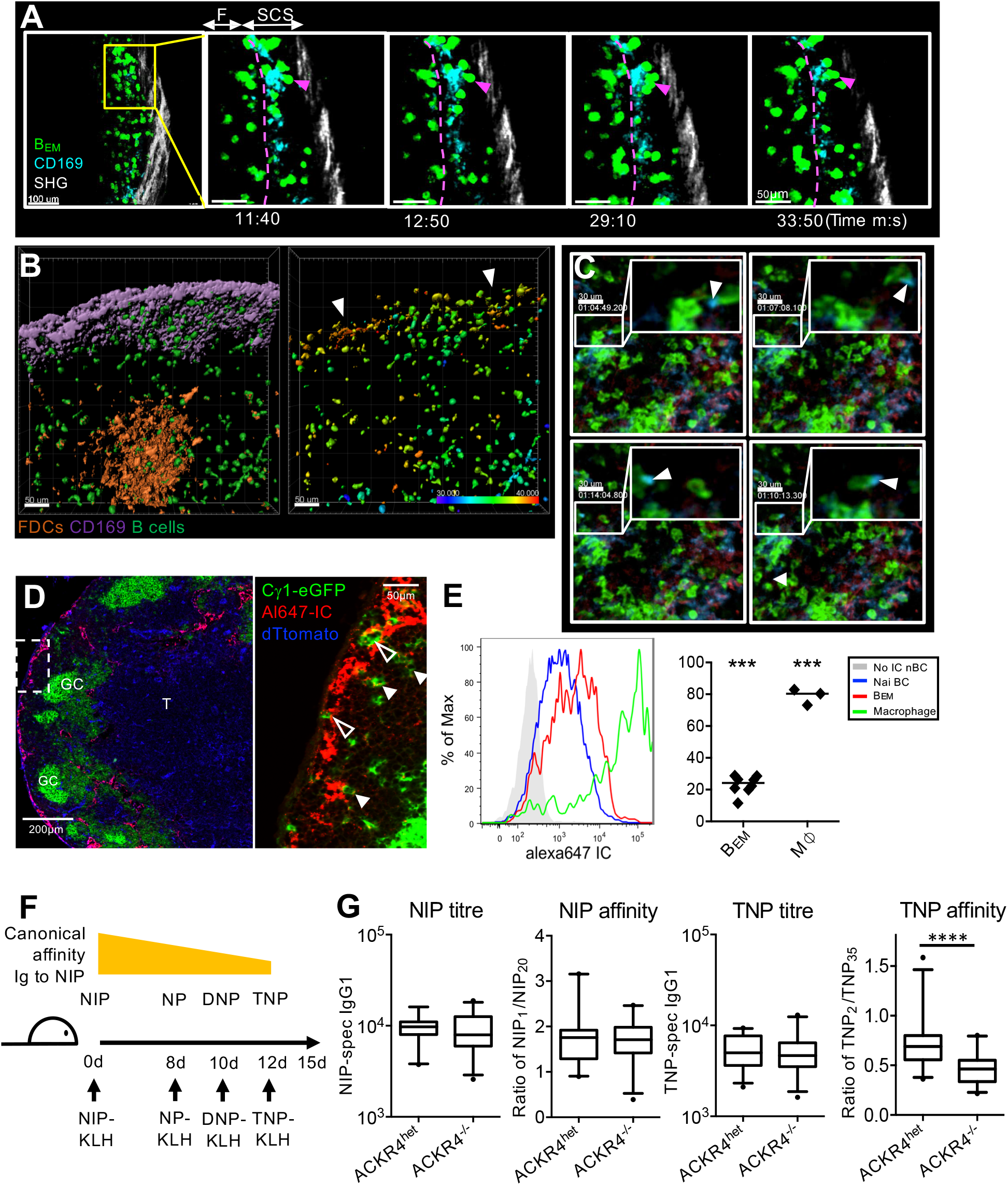
B_EM_ interaction with antigen in SCS affects response to antigenic variants. **A)** Intravital observation of interactions of eGFP^+^ B_EM_ (green) and SCS macrophages (CD169 turquois). **B**) Intravital Ca^2+^ levels in B_EM_. Left: Surface rendering of CD169-stained macrophages (purple) and GC (orange, CD21/35^Atto590^), B1-8^hi^ TN-XXL^+^ B cells (green). Right: FRET intensity in B_EM_ of same frame with color-coded mean FRET intensity. Scale bar 50 µm. **C**) Intravital observation (clockwise from top left) of B_EM_ (green) in SCS containing CD169^+^ material (blue). **D**) drLN from an Cγ1Cre mTmG mouse 10 min after foot injection with Alexa647 labelled immune complex (IC). Open Arrows: eGFP^+^ B_EM_ in SCS. Closed arrows: Alexa647-IC colocalizing with eGFP^+^ B_EM_ in follicle. **E**) Left: Alexa647-IC on CD169^+^ SCS macrophage (green), eGFP^+^ B_EM_ (red), naïve B cells (blue), or naïve B cells without Alexa647 injection (grey). Right: Percent Alexa647-IC positive B_EM_ or macrophages compared to naïve B cells. Overton subtraction from two independent experiments with 3 mice each. Unpaired T test,*** p<0.001. **F)** Design of antigenic drift experiment using variants of the NIP hapten. **G)** NIP-specific and TNP-specific IgG1 antibody titre and affinity in ACKR4^het^ and ACKR4^-/-^ mice. Box plots are merged data from 4 independent experiments, Mann-Whitney test, ****p<0.0001

The non-signaling atypical chemokine receptor 4 (ACKR4) is expressed in the SCS ceiling endothelium and shares the ligands CCL19 and CCL21 with CCR7. ACKR4 generates the CCL21 chemokine gradient that guides DCs into lymph nodes (Ulvmar et al., 2014). To test whether CCR7-mediated retention of B_EM_in the drLN is dependent on an ACKR4-generated chemokine gradient, we co-transferred QM mT^+^*Ackr4*^+/+^and QM eYFP^+^ *Ackr4*^-/-^ B cells into *Ackr4*^+/+^ or *Ackr4*^-/-^ hosts and immunized with NP-CGG. While ACKR4-deficiency on B cells had no significant effect on the size of the GC compartment nor affected B_EM_ numbers in the drLN, ACKR4-deficiency of the LN environment led to decreased numbers of antigen-specific B_EM_ being retained in the drLN and higher numbers appearing in the blood (Fig. 3D). This suggests that chemotactic cues generated in the SCS environment organize B_EM_ reentry into the drLN. In the absence of these, B_EM_ leave the SCS in larger numbers to appear as B_CM_ in the efferent lymph and distLNs.

To further test this, we followed the accumulation of B_EM_ in the SCS of drLN by fluorescence microscopy in immunized Cγ1Cre mTmG mice that were *Ackr4*^+/+^or *Ackr4*^-/-^. This revealed a significantly increased retention of B_EM_ in the SCS of ACKR4^-/-^ drLN 8 d when B_EM_ recycle in the drLN (Fig. 3E, F). There was also an increased number of B_CM_ arriving in ACKR4^-/-^ distLNs, and this difference persisted until d 14 (Fig. 5F). Intravital two-photon microscopy confirmed the increased numbers of B_EM_ in the SCS of drLNs of ACKR4-deficient mice (Fig 5G, H, suppl. Fig. S1, suppl. movie 3). Importantly, B_EM_ re-entry from the SCS into the lymph node was rarely observed when ACKR4 was absent (suppl. movie 4).

### B_EM_ recycling supports adaption to antigenic drift

We next considered the functional significance of B_EM_ LN re-entry from the SCS into the B cell follicle. Of note, we observed that B_EM_ appeared to undergo prolonged interaction with CD169-positive SCS macrophages (Fig. 4A, suppl. Fig. S2, suppl. movie 5) before reentering the LN parenchyma. Intravital imaging of cytoplasmic calcium levels showed an increase in calcium specific to B cells contacting SCS macrophages (Fig. 4B, suppl. Fig. S3), suggesting an antigen-specific interaction between B_EM_and antigen-carrying SCS macrophages. SCS macrophages are known to transfer antigens to naïve B cells (Carrasco and Batista, 2007; Junt et al., 2007). A recent study showed similar interactions of antigen-specific memory B cells during secondary responses (Moran et al., 2018). For some B_EM_, we observed that they acquired CD169-labelled material from SCS macrophages (Fig. 4C, suppl. movie 7), suggesting that B_EM_ acquire and transport antigen from SCS macrophages into the GC. To test this, mice were immunized with rabbit-IgG and eight days later injected with AlexaFluor647-labelled mouse anti-rabbit immune complex (IC). Within 10 min, IC was seen associated with SCS macrophages. IC was also present inside intranodal lymphatics and entering the lymph node parenchyma. B_EM_ within the SCS were in intimate contact with IC-carrying cells, whereas inside the LN parenchyma, those B_EM_ that were close to the SCS carried speckles of IC (Fig. 4D). Flow cytometry confirmed that 20 – 30 % of B_EM_ carried increased amounts of IC within minutes of IC injection (Fig. 4E). Together, these data suggest that B_EM_ may be activated by specific antigen in the SCS, and can transport this back into the LN parenchyma.

GCs typically contain large amounts of antigen deposited on follicular dendritic cells (FDC). Therefore, additional antigen deposition by B_EM_ seems unnecessary, unless the antigen is changing during the course of an infection. B_EM_, are a GC output with highly variable affinity and specificity for antigen, and would therefore include cells that may interact with antigenic variants. To test the hypothesis that B_EM_ recycling has a role in adaption to antigenic drift, we used variants of the hapten NP and measured the adaption of affinity maturation to these variants. The B cell response of C57BL/6 mice is dominated by a canonical IgH VDJ BCR combination that has natural affinity to 4-hydroxy-iodo-phenyl (NIP), and reduced affinity to the variants NP, dinitrophenyl (DNP) and trinitrophenyl (TNP) (Fig. 4F). C57BL/6 mice were immunized with NIP-KLH. After the onset of B_EM_ recycling, we rechallenged in the same foot with NP, DNP, followed by TNP-KLH. Three days after the last injection we observed a shift in antibody affinity towards TNP (suppl. Fig. S4). In order to test whether this was dependent on B_EM_ recycling, the experiment was repeated in Ackr4^ko^ mice, where B_EM_ cannot undergo recycling. This showed that without B_EM_ recycling, the drift towards the new antigenic variant was significantly reduced (Fig. 4G).

The SCS is the site of antigen entry into the lymph node. We have shown that B_EM_ collect antigen from this important anatomical site, as described for naïve B cells. For naïve B cells, however, this is mediated mainly by non-antigen-specific receptors (Carrasco and Batista, 2007). Many pathogens, particularly viruses, mutate frequently leading to antigenic drift that, during the course of a primary infection, can lead to a large accumulation of VDJ gene mutations (Schoofs et al., 2019). The recycling of highly variable B_EM_ through the SCS described here allows for BCR-dependent selection of B cell variants reacting with mutated antigen. This may accelerate affinity maturation to antigenic drift, not only because B_EM_ transport antigenic variants back into the GC, but more likely because they provide genomic code for immunoglobulin variants that can kickstart affinity maturation to the variant antigen. We do not know whether B_CM_ upon leaving the reactive lymph node act in a similar way when they encounter antigen in other sites. While some low affinity B cells can enter GCs during recall responses (Wong et al., 2020), most mature memory B cells are primed to differentiate into plasma cells (Mesin et al., 2020; Moran et al., 2018; Toellner et al., 1996; Viant et al., 2020). Whether this change in memory B cell function matures with time in absence of stimulation by antigen, or is induced by specific environments remains to be seen.

## Supporting information

Supplementary Mov1

Supplementary Mov2

Supplementary Mov3

Supplementary Mov4

Supplementary Mov5

Supplementary Mov6

Supplementary Mov7

## Abbreviations

B_EM_: memory like B cell emerging from germinal centers
B_CM_: circulating memory B cell
S1P: sphingosine-1-phosphate
GC: germinal center
ACKR4: atypical chemokine receptor 4
NP-CGG: 4-hydroxy-nitrophenyl coupled to chicken gamma globulin
QM: quasi-monoclonal mouse
SCS: subcapsular sinus
pLN: popliteal lymph node
drLN: draining lymph node
distLN: distant lymph node
DC: dendritic cell
FDC: Follicular dendritic cells
IC: immune complex

## Authors contributions

Conceptualization: YZ, AR, AEH, MRC, KT, Methodology: YZ, LGI, CU, MRC, Formal analysis: CJB, Investigation: YZ, LGI, CU, LSL, TWD, JRF, JMW, CJB, JYP, LZ, Writing: YZ, LGC, CU, MRC, KMT, Writing - review and editing: GB, VLT, AEH, MRC, KMT, Visualisation: CU, LSL, TWD, JRF, Supervision: YZ, VLT, AEH, MRC, KMT, Funding acquisition: YZ, LSL, GB, VLT, AEH, MRC, KMT All data is available in the manuscript or the supplementary materials.

## Supplementary Materials

Materials and Methods

Fig S1 – S4

References (1-16)

Movie 1-7

## Supplementary materials and figures

### Materials and Methods

#### Mice and immunization

C57BL/6J mice (wild type, WT) were purchased from Harlan laboratories. ACKR4^tm1.1Rjbn^ (ACKR4^-/-^) mice (1) were a gift from R. Nibbs (University of Glasgow). Cγ1Cre mTmG mice were generated by crossing Ighg1^tm1(cre)Cgn^ (*2*) with Gt(ROSA)26Sor^tm4(ACTB-tdTomato, -EGFP)Luo^ mice (*3*) (Jackson lab). For all adoptive transfer experiments, variants of QM mice were used, which were homozygous NP-specific Ig heavy chain variable region from Igh-J^tm1(VDJ-17.2.25)Wabl^ and Igk^tm1Dhu^ (*4*). Some QM mice contained a constitutively expressed enhanced yellow fluorescent protein (eYFP) derived from Gt(ROSA)26Sor^tm1.1(EYFP)Cos^ (*5*) (QM eYFP mice), if crossed with Rosa26mTmG, called QM mT. QM CCR7 mice are QM crossed with CCR7^tm1Rfor^ mice (*6*). Animal studies were performed with approval of local ethical committees and under appropriate governmental authority.

NP (4-hydroxy-3-nitrophenyl acetyl) was conjugated to CGG (Chicken γ-globulin) at a ratio of NP_18_-CGG. Mice were immunized into plantar surface of their rear feet with 20μg NP_18_-CGG alum precipitated plus 10^5^ chemically inactivated *Bordetella pertussis* (B.p.) (LEE laboratories, BC, USA) (*7*). Popliteal lymph node (pLN) were analyzed as reactive (or draining) lymph nodes, and axillary and brachial lymph nodes as remote (or distant) lymph nodes.

FTY720 (Caymanchem, USA) was given at 1mg/kg body weight via *i*.*p*. at d6 and d7 after NP-CGG on feet of mice which received 2×10^5^ NP^+^ B220^+^ cells. For immune complex injections, 4 μg of Alexafluor 647 labelled immune complex (IC) was made with 1:1 ratio of Alexafluor647 conjugated mouse anti-rabbit IgG plus rabbit IgG, mixed 30 minutes before injection into the foot, 8 d post priming with 20 μg rabbit IgG alum precipitated with 10^5^ B.p.

For antigenic drift experiments ACKR4^-/-^ mice and litter mate wild type control mice were primed with 10 μg of NIP-KLH in alum precipitated with 10^5^ B.p. in the rear feet. 8 d later, mice were boosted with 1 μg of soluble NP-KLH, DNP-KLH, and TNP-KLH on the same feet every 2 days (NP conjugates from Biosearch Technologies, USA).

#### Immunohistology

Lymph node sections were prepared and stained as described previously (*8, 9*). CD138 (281-2) and IgD APC (11-26c, BD BioSciences), biotinylated peanut agglutinin (PNA) (Vector Labs), Ki-67 and LYVE-1 (Abcam), ER-TR7 (eBioscience) were used. Secondary antibodies were Cy3-conjugated donkey anti-rat or donkey anti-rabbit (Jackson ImmunoResearch Laboratories, West Grove, PA) and Alexafluor 405 conjugated streptavidin (Invitrogen UK). Slides were mounted in ProLong Gold antifade reagent (Invitrogen, UK) and left to dry in a dark chamber for 24 h. Images were taken on a Leica DM6000 fluorescence-microscope, or Zeiss Axio ScanZ1. Image data were processed using Fiji (*10*) or ZEN (Carl Zeiss Germany).

#### Flow Cytometry and adoptive transfer

Cells from spleens and lymph nodes were prepared as described (*8*). Red blood cells were lysed by ACK lysing buffer (Gibco). Cell suspensions were blocked by CD16/32 (93, eBioscience) diluted in FACS buffer (PBS supplemented with 0.5% BSA plus 2mM EDTA), followed with staining cocktail. PD-L2 biotin (TY25) and B220-BV421 or BV510 (RA3-6B2) and Str. BV421 or BV711 were from BioLegend, NP was conjugated in house with PE for detected antigen specific B cells (*8*). CD38 APC (90), GL7 eFluor 450 (GL7), CD86 PE-Cy5 (GL1), CD80 PE-Cy5(16-10A1), and CD73 PE-Cy7 (eBioTY) were from eBioscience. CXCR4 biotin (2B11), Fas PE-Cy7 (Jo2) and CD138 APC or BV421 (281-2) were from BD Bioscience. Samples were analyzed using BD LSRFortessa Analyzer (BD Biosciences, USA) with the software BD FACSDiva (BD Biosciences). Data were analyzed offline with FlowJo (FlowJo LLC, USA). For adoptive transfer experiment, 2×10^5^ NP^+^ B220^+^ cells from spleens of fluorescent protein labelled QM background mice were transferred into C57BL6/J hosts 1 d before immunization with NP-CGG in alum on rear feet. In co-transfer experiments, a mix of 1×10^5^ of NP^+^B220^+^ B cells of each genotype respectively were injected *i*.*v*.

#### Intravital microscopy

Intravital microscopy of popliteal lymph nodes of Cγ1Cre mTmG mice were performed 8 days after immunization with NP-CGG in alum ppt on plantar surface of rear feet. Subcapsular sinus macrophages were labelled with CD169-A647 antibody (BioLegend) injected subcutaneously to foot before imaging. Popliteal LN imaged under anesthesia with a Chameleon Ti:Sapphire multiphoton laser and Leica SP8 microscope. Images were acquired using a 25x objective, with one Z stack every 30 to 40 seconds, and processed using either Bitplane Imaris or Fiji ImageJ (*11*).

#### Cell sorting for qRT-PCR, RNA-seq library preparation and data analysis

Draining LN and distant LN in mice 8 d after foot immunization with NP-CGG in alum and B.p were stained as described above. Naive B cells, GC B cells, plasma cells, B_EM_ cells from drLN and B_CM_ from distLN were sorted using a high speed cell sorter (MoFlo, Beckman-Coulter).

For real time PCR, RNA was purified by using the RNeasy Mini kit (QIAGEN), cDNA preparation was as described as before (*8*). Real-time PCR from cDNA (qRT-PCR) was done in multiplex with β2-microglobulin and gene expression related to β2-microglobulin gene expression levels. Primers and probes are listed in Zhang et al 2018 (*12*), and as follows: S1pr2 Fwd: GGCCTAGCCAGTGCTCAGC, Rev: CCTTGGTGTAATTGTAGT, probe: FAM-CAGAGTACCTCAATCCTGA-TAMRA. CCR7 Fwd: GGTGGCTCTCCTTGTCATTTTC, Rev: GTGGTATTCTCGCCGATGTAGTC, probe: FAM-TGCTTCTGCCAAGATGAGGTCACCG-BHQ1. Cxcr3 (Mm00438259_m1), Ccr6 (Mm01700300_g1), Ebi2 (Mm02620906_s1) were TaqMan gene expression assays (Thermo Fisher Scientific, UK). S1pr1 and Ackr4 were run with SYBRGreen realtime PCR (Thermo Fisher Scientific). S1pr1, Fwd: AAATGCCCCAACGGAGACT, Rev: CTGATTTGCTGCGGCTAAATTC. Ackr4: Fwd: TGG ATC CAA GAT AAA GGC GGG GTG T143YES, Rev: TGA CTG GTT CAG CTC CAG AGC CAT G For RNA-seq, cells were directly sorted into 500ul of Trizol. The total RNA was purified using the RNeasy Plus Micro kit (QIAGEN) according to the manufacturer’s instructions. Un-stranded, non-rRNA, non-polyA+ selected libraries were prepared using the SMARTer Ultra Low Input RNA kit for Sequencing v3 (Clontech Laboratories). The libraries were sequenced on the Illumina HiSeq 2000 platform (Illumina, Crick advanced sequencing) as 75 bp paired-end runs.

The sequencing data was analyzed using Partek^®^ Flow^®^ software, version 8.0.19 Copyright ©; 2019 Partek Inc., St. Louis, MO, USA. Paired sequencing data was imported and then aligned to mouse genome GRCm38 (mm10). t-SNE analysis was performed on normalized RNA counts to generate a 2D plot by dimensional reduction. Gene specific analysis (GSA) tool was used to identify differentially expressed genes against naïve B cells subset as a control. GSA used the lognormal and negative binomial response distribution under the multi-model approach and a lowest maximum coverage of 1.0 was used as the low-value filter. Venn diagrams were produced from differential expression of genes with a log fold change >1 and P-value <0.05 using BioVenn (Hulsen et al., 2008). The heatmap was produced from GSA data as described earlier from a predetermined list of genes of interest using the Hierarchical Clustering tool; genes were clustered based on their average Euclidean distance from one another.

#### Calcium imaging by intravital microscopy

C57BL6/J mice received B cells from B18hi mice (carrying the Vh186.2 heavy chain with high affinity to NP) that contained genetically encoded Ca^2+^ indicator TN-XXL (*13*) under the control of the CD19 promoter (*14*). These were immunized with 10 µg NP-CGG emulsified in complete Freund’s adjuvant (CFA) into the right foot. The popliteal LN was analyzed at day 7. One day prior to imaging, a mixture of 10 µg anti-CD21/35 Fab (clone 7G6) -Atto590 (produced at the DRFZ) for staining follicular dendritic cells and 10 µg CD169-efluor660 (eBioscience) to label SCS macrophages were injected into the footpad.

Intravital two-photon microscopy was performed as described before (*15*), using a TrimScope II from Lavision Biotec, at an excitation of 850 nm (TiSa) and 1100 nm (OPO). The detection of the fluorescence signals was accomplished with photomultiplier tubes in the ranges of (466 ± 20) nm, (525 ± 25) nm and (655 ± 20) nm.

TN-XXL is a genetically encoded calcium indicator that consists of a chicken troponin C domain connecting the fluorescent proteins eCFP and Citrine (Suppl. Fig. S7A). These act as a Förster resonance energy transfer (FRET) pair with ECFP as the donor and Citrine as the acceptor fluorophore. Troponin C contains four binding lobes for Ca^2+^ ions. If Ca^2+^ is present or cytosolic concentrations are elevated this leads to a conformation change of the linker peptide that causes donor and acceptor to come into sufficient proximity for FRET emission. When quenched ECFP is excited with one photon at 475 nm, or two photons at 850 nm, citrine will emit fluorescence at 530nm. If no calcium is present, emission in the blue range of the donor group will be more prominent.

Measurements from six immunized mice were analyzed with image analysis software Imaris (Bitplane AG). Raw data was pre-processed using a linear unmixing algorithm (*16*) to minimize interference of red fluorescence from antibody staining into the green channel of the citrine fluorescence. Relative FRET ratio was calculated from dividing green fluorescence gain by the sum of blue and green fluorescence, and corrected for instrument-specific values and spectral overlap. A colocalization channel was used to measure contact intensity between B cells (citrine-positive, masked on eCFP to exclude OPO influence) and CD169-efluor660 signal. Using the histogram of the colocalization intensity mean of the B cell surfaces, we identified distinguishable populations of B cells (Suppl. Fig. S7F) with either no contact (-) or tight contact (+). All B cells with colocalization intensity of 0 AU were assigned to the (-) group. To choose a threshold value of colocalization intensity for B cells to be assigned to the (+) group, we biexponentially fitted the decay of the histogram and determined the point in which cell numbers intersect the plateau of y=9,509 to be 717 AU. Non-contacting B cells and B cells with colocalization intensities >717 AU were filtered and corresponding FRET intensities of all cells at all time points exported for plotting.

#### Statistical analysis

All analysis was performed using GraphPad Prism 6 software. To calculate significance two-tailed Mann-Whitney non-parametric test was used. In the experiments where 2 parameters from the same individual mouse are compared, Wilcoxon matched-pairs signed rank test (paired non-parametric test) was used to calculate significance. Statistics throughout were performed by comparing data obtained from all independent experiments. P values <0.05 were considered significant (*). *p<0.05, ** p< 0.01, *** p<0.001, ****p<0.0001

## Supplementary figures

**Supplementary Fig S1:**
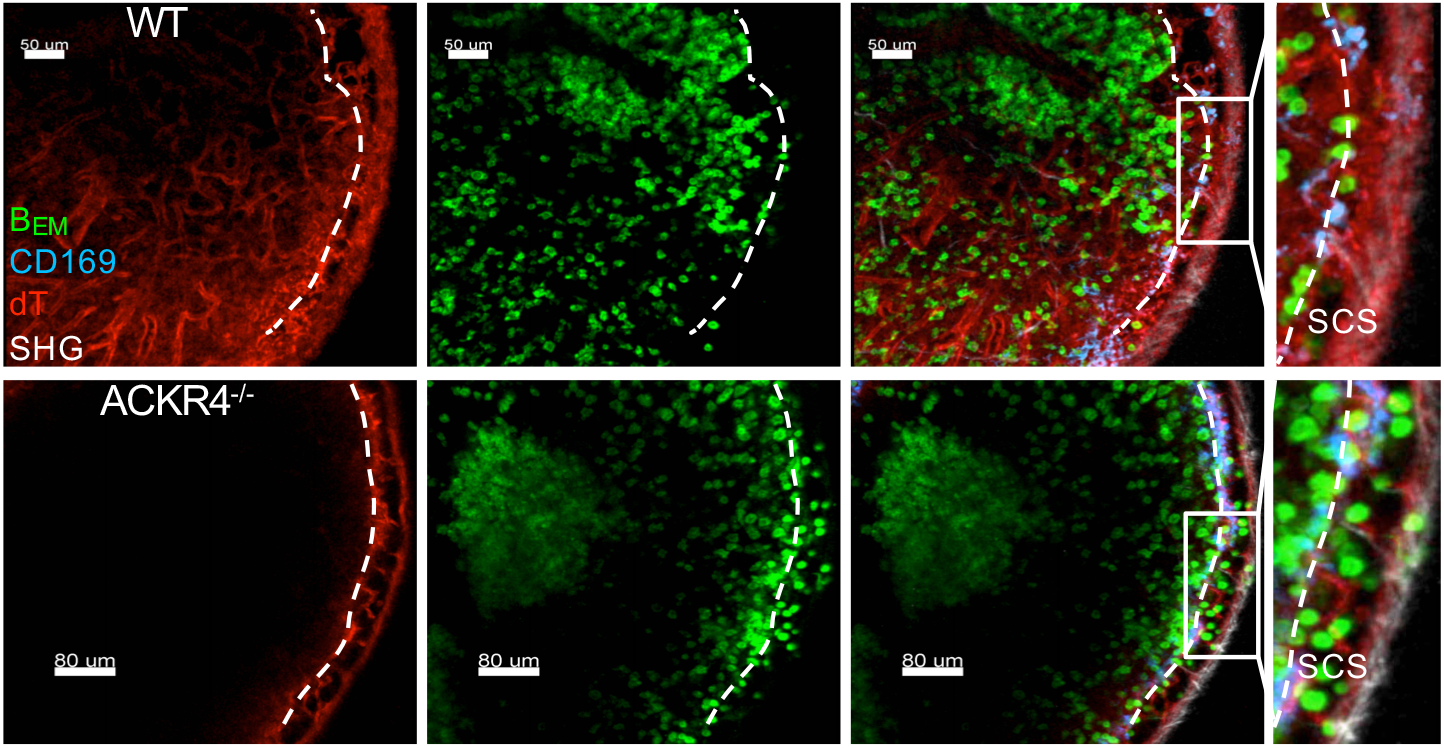
more B_EM_ in SCS of ACKR4 deficient drLN. Still images of intravital microscopy of Cγ1Cre mTmG ACKR4^+/+^ (top) and ACKR4^-/-^ (bottom) in drLN 8 d after foot immunization. This shows eGFP-labelled B_EM_ (green), mTomato-labelled stroma (red) and CD169 labelled SCS macrophages (blue). Note larger numbers of B_EM_ in the SCS in ACKR4^-/-^ environment.

**Supplementary Fig S2:**
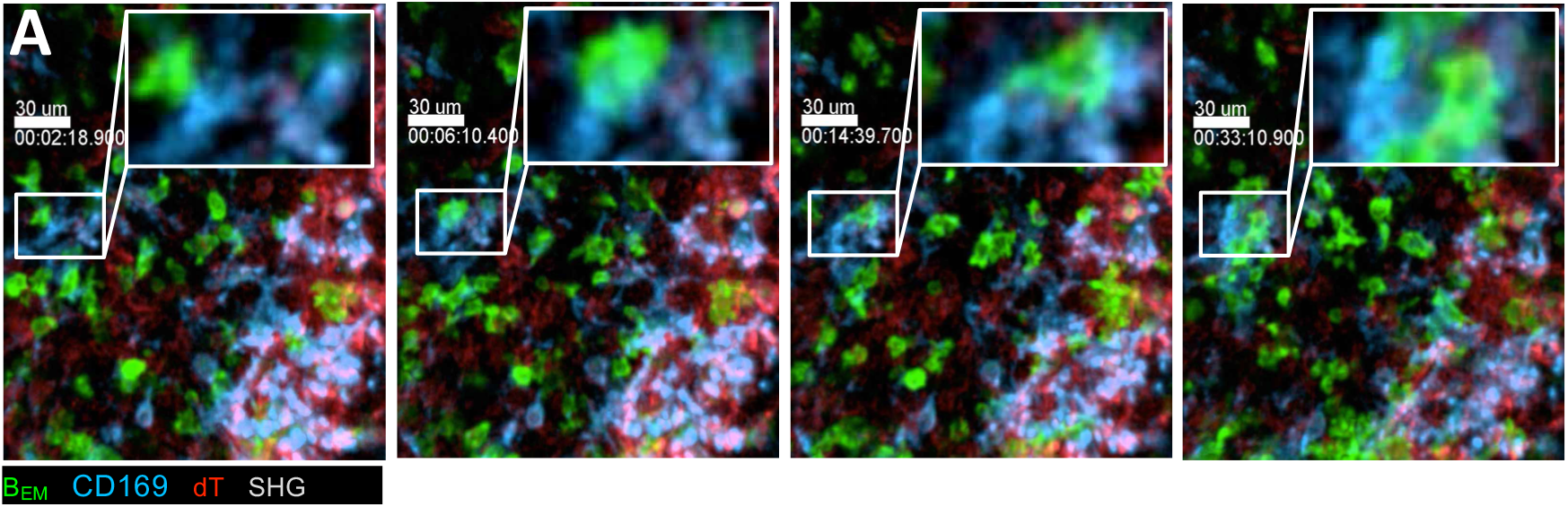
Intravital observation of B_EM_ interacting with CD169^+^ SCS macrophage. Still images from intravital microscopy of SCS in Cγ1Cre mT/mG ACKR4^+/+^ drLN 8 d after foot immunization. eGFP^+^ B_EM_ make prolonged interaction with CD169-labelled SCS macrophages (blue). mTomato positive stroma (red).

**Supplementary Fig S3:**
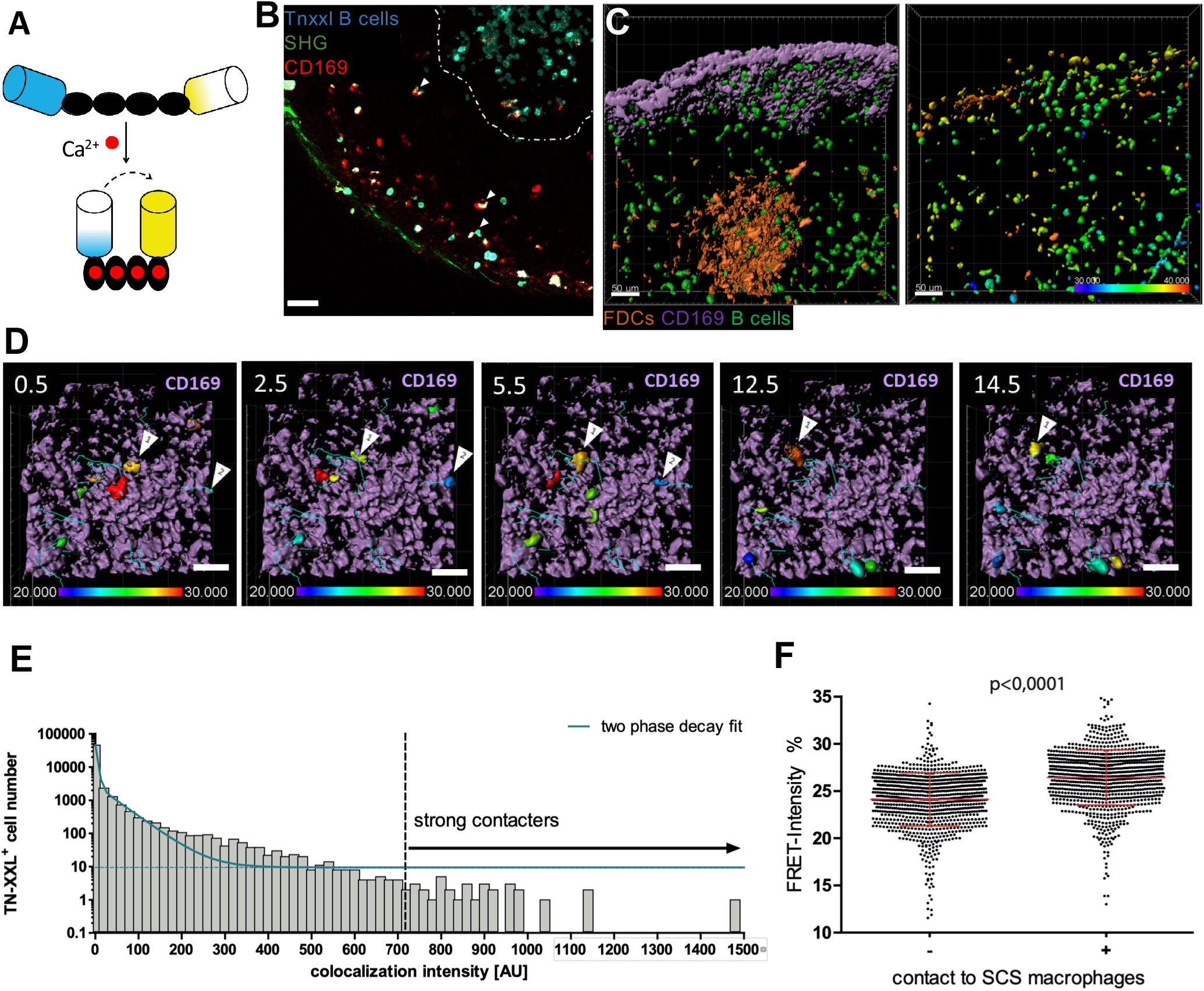
Intravital observation of intracellular Ca^2+^ in B_EM_. **A)** Schematic representation of the genetically encoded calcium indicator TN-XXL. The four troponin c binding sites of TN-XXL (black ovals) can be loaded with four calcium ions (red) that cause a conformational change. Förster resonance energy transfer leads to decreased eCFP (blue) and increased citrine (yellow) fluorescence after multiphoton excitation with 850nm laser light. **B)** Single z plane of intravitally imaged lymph node with capsule visible (SHG), CD169-efluor660 stained macrophages (red) and adoptively transferred TN-XXL+ B1-8^hi^ (NP-specific) B cells (cyan). A germinal center is visible at the upper right (broken white line). Contacts between CD169+ macrophages and TN-XXL^+^ B cells are shown in white (white arrows). **C)** Left: Surface rendering of subcapsular sinus (light purple, CD169-stained macrophages) and germinal center (light orange, FDC staining with CD21/35-Atto590), TN-XXL^+^ B1-8^hi^ B cells (green). Right: FRET intensity of TN-XXL^+^ B cells shown in left image, color-coded for mean FRET intensity. Scale bar 50 µm. See also suppl. movie S6. **D)** Stills of TN-XXL^+^ cells tracked over 30 minutes, mean FRET values depicted as color coding. CD169+ macrophages in light purple, track lines in cyan. White arrows point to tracks analyzed in E) and F) for FRET and colocalization intensity over time. Scale bar 40 µm. **E)** Colocalization histogram and two-phase decay fit for analysis of colocalization between CD169^+^ macrophages and TN-XXL^+^ B cells. The colocalization intensity where the decay reaches a plateau was determined as threshold for strong (+) macrophage-to-B-cell contact. B cells with colocalization intensity of 0 AU were assigned to the (-) group. **F)** Exemplary plot of mean FRET intensities of surface rendered B cells gated on contact strengths to SCS macrophages: (-) group: colocalization intensity = 0 AU; (+) group: colocalization intensity > 717 AU. One dot per TN-XXL+ object. Plot is representative for 5 individual experiments. Students t-test with Whelch correction.

**Supplementary Fig S4:**
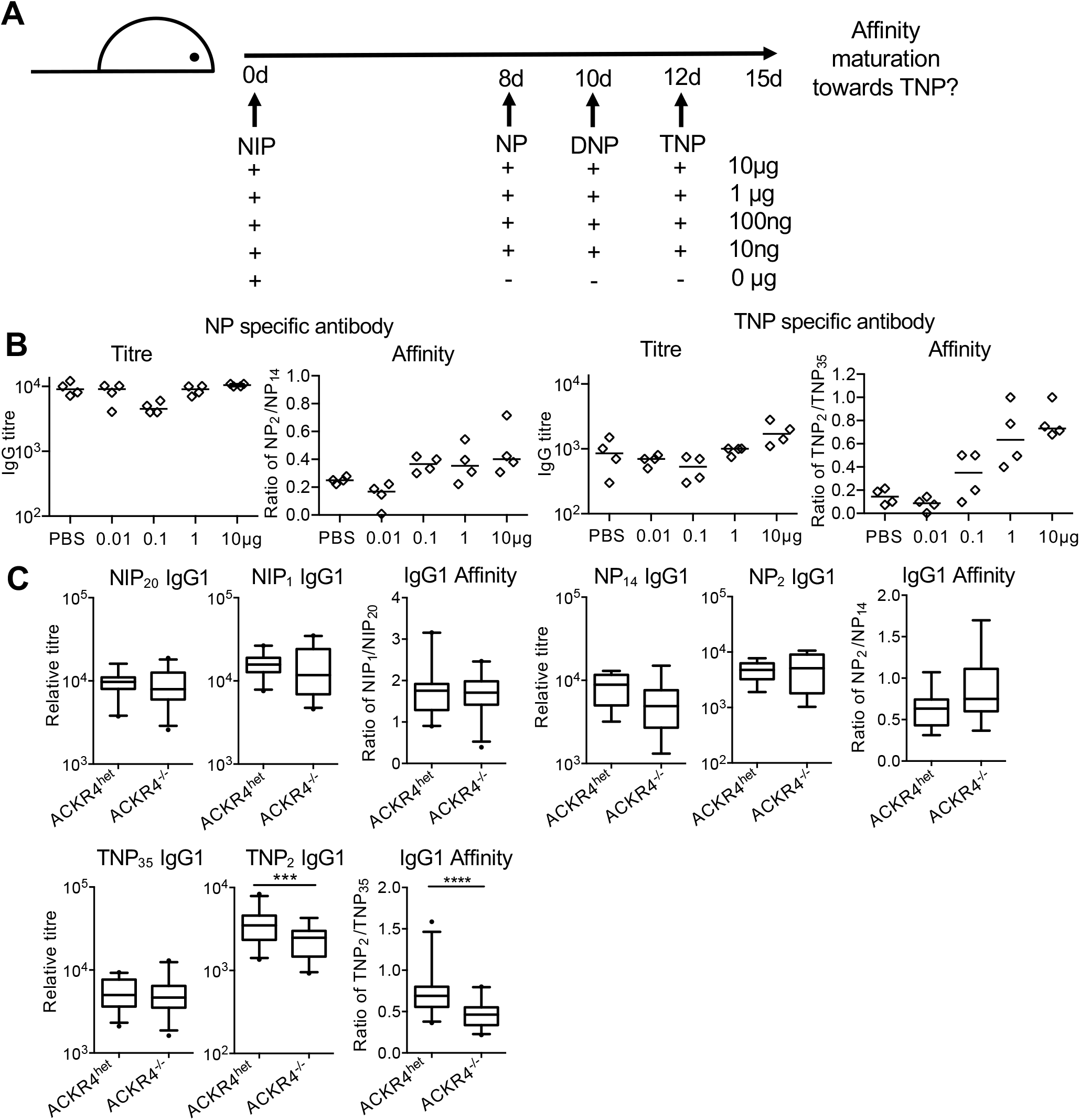
Antigenic drift experiment. **A)** Experimental design to simulate antigenic drift. Wild type mice were primed with 20 µg / 20 µl NIP-KLH in alum with B. pertussis into foot, followed by subsequent injections 8, 10, and 12 d after immunization with soluble NP-, DNP-, TNP-KLH into the same foot respectively. Control mice received mock immunization. Blood was collected 15 days post-immunization. **B)** NP- or TNP-specific antibody titer and affinity. Each dot represents one animal. ANOVA with Tukey test for multiple comparisons, * p<0.05, ** p<0.01, *** p<0.001, **** p<0.0001. **C)** NIP-, NP-, or TNP-specific antibody titers and affinity after immunization of ACKR4^wt^ or ACKR4^ko^ litter mates with NIP-KLH followed by NP-, DNP-, TNP-KLH. Merged data from four independent experiments with 5-6 mice.

## Supplementary movies

**Supplementary movie 1: movement of B**_**EM**_ **between GC and SCS and in and out of the SCS.**

Intravital microscopy. Overview of a Cγ1Cre mTmG ACKR4^+/+^ drLN day 8 after foot immunization. Cγ1Cre-dependent expression of eGFP (green) shows GC and B_EM_. CD169 (blue) indicates location of SCS with SCS macrophages. Red: dTomato expressing stroma. Grey: second harmonic.

**Supplementary movie 2: GFP**^**+**^ **B**_**EM**_ **moving into the SCS and reentry into drLN**

Intravital microscopy. Higher power video of SCS of a Cγ1Cre mTmG ACKR4^+/+^ drLN 8 d after foot immunization. Cγ1Cre-dependent expression of eGFP (green) shows GC and B_EM_. CD169 (blue) indicates SCS macrophages. Red: dTomato expressing stroma. Grey: second harmonic. Blue line: Track of B_EM_ entering the SCS. White line: Track of B_EM_ returning into the lymph node from the SCS.

**Supplementary movie 3: Location of B**_**EM**_ **in relation to SCS macrophages and SCS lumen in wt and ACKR4**^**ko**^ **drLN**.

Intravital microscopy. 3D still image of SCS of a Cγ1Cre mTmG ACKR4^+/+^ drLN 8 d after foot immunization. Cγ1Cre-dependent expression of eGFP (green) indicating B_EM_. CD169 (blue) SCS macrophages lining the SCS floor endothelium. Red: dTomato expressing stroma. Grey: second harmonic indicating the LN capsule. B_EM_ can be seen inside the LN parenchyma and having entered the SCS.

**Supplementary movie 4: B**_**EM**_ **moving along the SCS in ACKR4**^**-/-**^ **drLN**

Intravital microscopy of SCS of a Cγ1Cre mTmG ACKR4^-/-^ drLN 8 d after foot immunization. Cγ1Cre-dependent expression of eGFP positive B cells (green and white). CD169 (blue) SCS macrophages lining the SCS floor endothelium. Red: dTomato expressing stroma. Tracked cells (white) can be seen moving inside the SCS, but not reentering the LN parenchyma.

**Supplementary movie 5: Prolonged interaction of B**_**EM**_ **with SCS macrophage**

Intravital microscopy of a Cγ1Cre mTmG drLN 8 d after foot immunization. Cγ1Cre-dependent expression of eGFP (green) indicating B_EM_. Red: dTomato expressing stroma. Blue: CD169 on SCS macrophages.

**Supplementary movie 6: Colocalization of B**_**EM**_ **with increased Ca2+ and SCS macrophages**

Animated GIF merging images from Fig. 6B, showing surface rendering of CD169^+ve^ macrophages (purple), FDC in GC (orange), GC B cells and B_EM_ (green), and on second frame FRET intensity of green B cells shown in first frame, color-coded for mean FRET intensity

**Supplementary movie 7: B**_**EM**_ **removing CD169**^**+**^ **material from SCS macrophage**

Intravital microscopy of a Cγ1Cre mTmG drLN 8 d after foot immunization. Cγ1Cre-dependent expression of eGFP (green) indicating B_EM_. Red: dTomato expressing stroma. CD169^+^ material (blue) can be seen at the trailing edge of the migrating B_EM_.

