## Supplementary figures and images for "Memory-like B cells emerging from germinal centres recycle through the subcapsular sinus"

### Supplementary Mov6

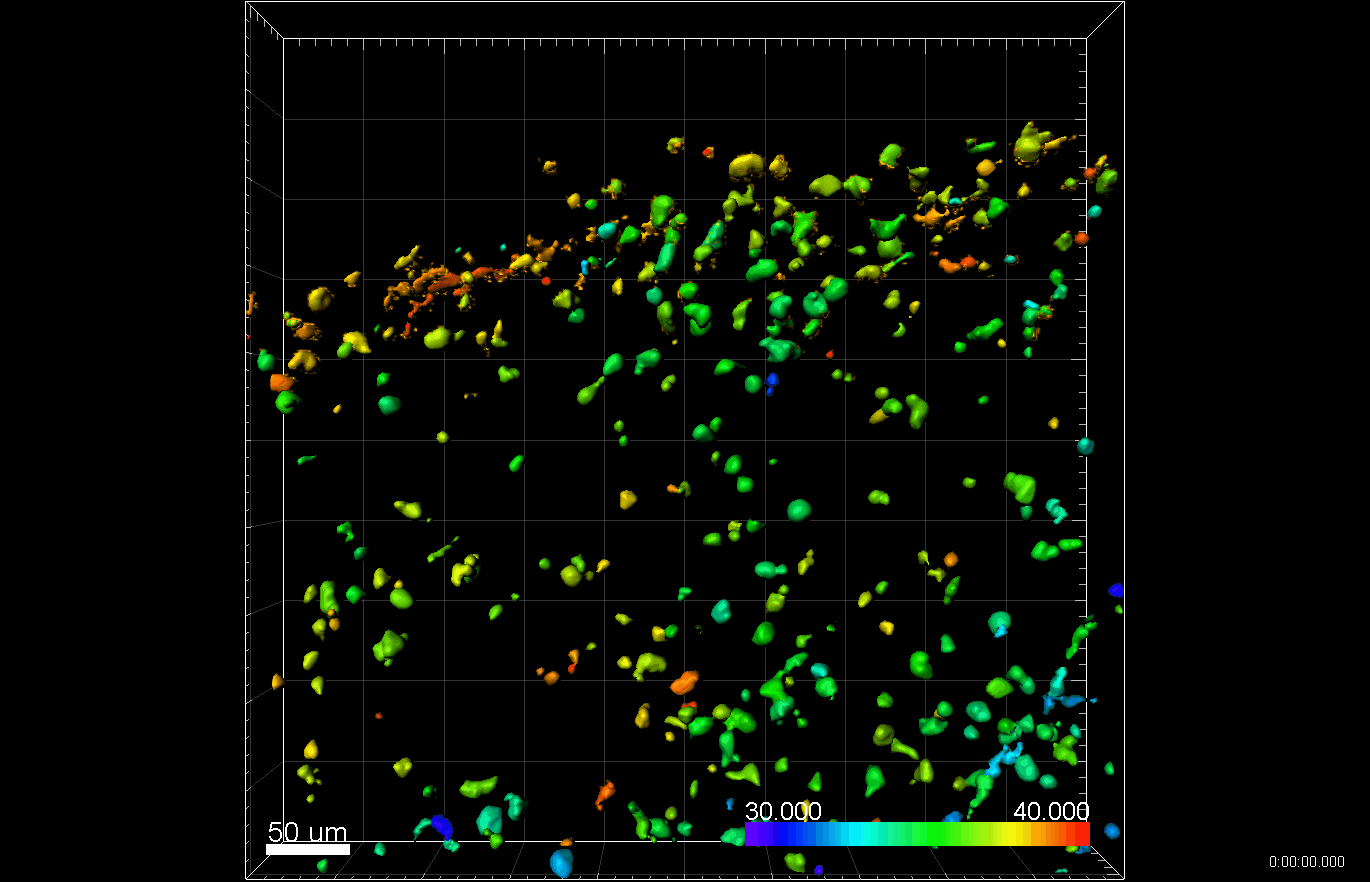
